# Desiccation Does Not Drastically Increase the Accessibility of Foreign DNA to the Nuclear Genomes: Evidence from the Frequency of Endosymbiotic DNA Transfer

**DOI:** 10.1101/465443

**Authors:** Xi-Xi Li, Cheng Fang, Jun-Peng Zhao, Xiao-Yu Zhou, Zhihua Ni, Deng-Ke Niu

**Affiliations:** MOE Key Laboratory for Biodiversity Science and Ecological Engineering and Beijing Key Laboratory of Gene Resource and Molecular Development, College of Life Sciences, Beijing Normal University, Beijing 100875, China

**Keywords:** Horizontal gene transfer, nuclear mitochondrial DNA (NUMT), nuclear plastid DNA (NUPT), double-strand breaks (DSBs), non-homologous end joining (NHEJ), bdelloid rotifers

## Abstract

Horizontal gene transfer (HGT) is a widely accepted force in the evolution of prokaryotic genomes. However, in eukaryotes, it is still in hot debate. Some bdelloid rotifers that are resistant to extreme desiccation and radiation were reported to have a very high level of HGTs. However, a similar report in another resistant invertebrate, tardigrades, has been mired in controversy. The DNA double-strand breaks (DSBs) induced by prolonged desiccation have been postulated to open the gateway of nuclear genome for foreign DNA integration and thus facilitate the HGT process. If so, the rate of endosymbiotic DNA transfer should also be enhanced. We first surveyed the abundance of nuclear mitochondrial DNAs (NUMTs) and nuclear plastid DNAs (NUPTs) in three groups of eukaryotes that are extremely resistant to desiccation, bdelloid rotifers, *A. vaga* and *A. ricciae*, tardigrades, *H. dujardini* and *R. varieornatus*, and the resurrection plants, *D. hygrometricum* and *S. tamariscina*. Excessive NUMTs or NUPTs have not been detected. Furthermore, we compared nine groups of desiccation-tolerant organisms with their desiccation-sensitive relatives but did not find significant difference in the NUMT/NUPT contents. Desiccation could induce DSBs, but it unlikely dramatically increase the frequency of foreign sequence integration in most eukaryotes. Only in the nuclear genomes enriched in repetitive sequences, the DSBs are predominantly repaired by non-homologous end joining (NHEJ) and desiccation-induced DSBs is possible to enhance the integration of foreign sequences into nuclear genome for some degree.

## Introduction

Horizontal gene transfer (HGT, also termed as lateral gene transfer) is the movement of genetic materials beyond the direction from parents to offspring (Keeling and Palmer 2008; Soucy, et al. 2015). It is well documented as the principal route of the evolutionary innovations in bacteria and archaea, like the acquirement of antibiotic resistance in pathogenic bacteria (Koonin 2016). By contrast, the contribution of HGT to the evolution of eukaryotic genomes is still in hot debates (Danchin 2016; Ku and Martin 2016; Martin 2017; McInerney 2017; Moreira and López-García 2017; Leger, et al. 2018). An example of the controversy is the amount of human genes acquired by HGT (Sieber, et al. 2017). In the first draft of the human genome, 223 proteins were found to be more similar with bacterial proteins than any eukaryotic protein sequences available at that time. These human proteins were claimed to arise from HGTs (Lander, et al. 2001). This result was quickly refuted by extensive analyses of the candidate genes (Salzberg, et al. 2001; Stanhope, et al. 2001). According to the opponents, gene loss in other eukaryotic lineages is the most likely explanation. 14 years later, Crisp et al (2015) reported that they confirmed 17 previous reported foreign genes and found 128 additional foreign genes in human genome in a large-scale comparative study. However, their conclusion was against by a case by case re-analysis of the 145 genes (Salzberg 2017). Almost at the same time, another group claimed that they discovered 1467 HGT regions in human genome, involving 642 known genes (Huang, et al. 2017). In eukaryotes, the most widely observed and well accepted HGT events are the transfer of organelle genome sequences to nuclear genomes, specially termed as endosymbiotic gene transfer (Timmis, et al. 2004; Kleine, et al. 2009; Ku, et al. 2015; Soucy, et al. 2015). After the endosymbiotic origin of mitochondria and plastids, most of their genetic sequences had been transferred to the nuclear genomes. Although effective transfer of functional protein-coding genes has slowed down or even stopped in some lineages, the process of DNA fragment transfer from organelle to nucleus is still actively continuing in most eukaryotic lineages (Timmis, et al. 2004; Hazkani-Covo, et al. 2010; Hazkani-Covo and Martin 2017a). The mitochondrial DNA segments recently inserted into nuclear genomes are termed as the nuclear mitochondrial DNAs (NUMTs), and the plastid DNA segments recently inserted into the nuclear genomes are termed as the nuclear plastid DNAs (NUPTs).

In most multicellular animals and plants, the opportunity of passing genetic information directly to future generations is restricted within a small proportion of cells, the germ line cells (Jensen, et al. 2016). This may serve as a strong barrier to HGT. But in both animals and plants there are a few weakly protected unicellular or early developmental stages in which there are some chances for environmental DNA to enter the germ cells and be inherited (Huang 2013). A striking animal in the field of HGT is bdelloid rotifers. Although the frequency of HGTs depends on the methods, the quality of the sequenced genome, and the specific species studied, a pattern of massive HGTs in bdelloid rotifers has been repeatedly reported (Gladyshev, et al. 2008; Boschetti, et al. 2012; Flot, et al. 2013; Eyres, et al. 2015; Nowell, et al. 2018). The HGTs in bdelloid rotifers are so frequent that some researchers suggest that it can act as an alternate form of sex to facilitate genetic exchange among bdelloid rotifers (Debortoli, et al. 2016; Schwander 2016). Although the evidence of HGT among bdelloid rotifers has been refuted in a most recent paper (Wilson, et al. 2018), so far as we know, there are no controversies on the massive HGTs from non-metazoan species to bdelloid rotifers. These tiny invertebrates live in ephemeral aquatic habitats and therefore occasionally experience severe desiccation. When facing desiccation, they enter a form of dormancy called anhydrobiosis (Gusev and Okuda 2012; Ricci 2017). It has been demonstrated that the prolonged desiccation of anhydrobiotic bdelloid rotifers induces DNA double-strand breaks (DSBs) (Hespeels, et al. 2014). The DSBs were vividly described as the “gateway to genetic exchange” by Hespeels et al. (2014) and are widely believed as the main cause of the elevated HGT frequency in bdelloid rotifers (Gladyshev, et al. 2008; Boschetti, et al. 2012; Flot, et al. 2013; Eyres, et al. 2015; Debortoli, et al. 2016).

Besides bdelloid rotifers, a number of other organisms also enter the anhydrobiotic state when their environments become drying. Another remarkable example is tardigrades (Gusev and Okuda 2012). Prolonged desiccation has also been shown to induce DSBs in tardigrades (Neumann, et al. 2009). By analogy with bdelloid rotifers, it is very natural to expect frequent HGT in tardigrades by a similar mechanism. However, tardigrades did not provide further support for the gateway hypothesis, but instead amplified the controversy of HGT in eukaryotes. The first analysis of a tardigrade draft genome (*Hypsibius dujardini*) suggested more extensive HGTs than bdelloid rotifers, 17.5% of its genes were identified as foreign genes originated from bacteria, plants, fungi, and archaea (Boothby, et al. 2015). However, subsequent independent sequencing of the same species as well as another tardigrade species, *Ramazzottius varieornatus*, consistently showed that the percentage of foreign genes in tardigrades is much lower, with 2.3% as the reported upper bound (Arakawa 2016; Bemm, et al. 2016; Delmont and Eren 2016; Hashimoto, et al. 2016; Koutsovoulos, et al. 2016; Yoshida, et al. 2017). Most previously identified foreign genes should result from contamination of DNA from non-target organisms. Accordingly, tardigrades should not be regarded as a special group of animals with elevated levels of HGT (Yoshida, et al. 2017). The predominant objections to the claimed extensive HGTs in tardigrades raise our suspicion on the gateway hypothesis. Could prolonged desiccation really open the “gateway to genetic exchange” by inducing DSBs in the nuclear genome? In this study, we hope to gain new insights on this question by analyzing the abundance of NUMT and NUPT of eukaryotic organisms that frequently experience desiccation. The rationale is that the hypothetical gateway opened by desiccation facilitates not only HGTs but also endosymbiotic gene transfers.

## Material and Methods

### Genomes of Desiccation-Tolerant Species and Their Controls

We downloaded the nuclear genome data of *Adineta ricciae* (GCA_900240375.1), *Ancylostoma duodenale* (GCA_000816745.1), *Apis mellifera* (GCF_003254395.2), *Chenopodium quinoa* (GCF_001683475.1), *Dorcoceras hygrometricum* (GCA_001598015.1), *Fagopyrum esculentum* (GCA_001661195.1), *Hypsibius dujardini* (GCA_002082055.1), *Lolium perenne* (GCA_001735685.1), *Mentha longifolia* (GCA_001642375.1), *Nilaparvata lugens* (GCF_000757685.1), *Ramazzottius varieornatus* (GCA_001949185.1), *Rotaria magnacaicarata* (GCA_900239745.1), *R. macrura* (GCA_900239685.1), *Selaginella moellendorffii* (GCF_000143415.4), *S. tamariscina* (GCA_003024785.1), and *Xeromyces bisporus* (GCA_900006255.1) from ftp://ftp.ncbi.nlm.nih.gov/genomes/ and those of *Adineta vaga* (AMS_PRJEB1171_v1), *Caenorhabditis briggsae* (CB4), *C. elegans* (WBcel235) *Daphnia magna* (daphmag2.4), *D. pulex* (V1.0), *Hordeum vulgare* (IBSC_v2), and *Paracoccidioides brasiliensis* (Paracocci_br_Pb18_V2) from ftp://ftp.ensemblgenomes.org/pub/release-41. The organelle genome data of *C. elegans* and *H. vulgare* were downloaded from ftp://ftp.ensemblgenomes.org/pub/release-41. The mitochondrial genome sequences of *A. duodenale* (NC_003415.1), *C. briggsae* (NC_009885.1), *D. magna* (NC_026914.1), *D. pulex* (NC_000844.1), *H. dujardini* (NC_014848.1), *L. perenne* (JX999996.1), *N. lugens* (NC_021748.1), *R. varieornatus* (NC_031407.1), *X. bisporus* (HG983520.1), and *P. brasiliensis* (NC_007935.1) and the plastid genome sequences of *D. hygrometricum* (NC_016468.1), *F. esculentum* (NC_010776.1), *L. perenne* (NC_009950.1), *M. longifolia* (NC_032054.1), and *S. moellendorffii* (NC_013086.1) were downloaded from https://www.ncbi.nlm.nih.gov/nucleotide/. The mitochondrial genome sequences of *A. vaga*, *A. ricciae*, *R. magnacaicarata*, and *R. macrura* were provided by the authors of the reference (Nowell, et al. 2018) and the plastid genome sequence of *S. tamariscina* was provided by the authors of the reference (Xu, et al. 2018). The annotation files of most organelle genomes were obtained along with the sequence data except those of the mitochondrial genomes of *A. vaga*, *A. ricciae*, *R. magnacaicarata*, *R. macrura*, and *X. bisporus*. We annotated these five genomes using MITOS WebServer (Bernt, et al. 2013).

### Detection and Filtration of NUMTs and NUPTs

The NUMTs and NUPTs of some model species have been explored for many times (Richly and Leister 2004a, b; Hazkani-Covo, et al. 2010; Smith, et al. 2011; Hazkani-Covo and Martin 2017a). Because different parameters used in previous studies and different levels of genome assembly would lead to discrepancies in NUPT/NUMT abundance (Smith, et al. 2011; Hazkani-Covo and Martin 2017a; Hazkani-Covo and Martin 2017b), we performed our own detection using the same set of parameters with the expectation that over- or underestimation affects all species consistently. The organelle genome sequences were used as the query to search against the nuclear genomes by blastn (version 2.4.0+) with a threshold E value of < 0.0001, a word size of 11, a match score of 2, a mismatch score of −3, and gap cost values of 5 (existence) and 2 (extension) as those used by Smith et al. (2011). To reduce false-positive hits resulting from contamination of organelle sequences in the nuclear genome sequences, short nuclear contigs that were completely matched to organelle genome sequences were discarded. Multiple organelle DNA segments that match the same nuclear DNA region were counted only once.

We also separately detected the NUMTs and NUPTs contributed by organelle sequences with known functions, including the protein-coding genes, rRNA genes, and tRNA genes using the above method and parameters.

## Results

### The Most Desiccation-Tolerant Animals and Plants Do Not Have Excessive NUMTs/NUPTs

Water deficiency is one of the most common abiotic stress factor for organisms living on land. To cope with environmental drying, terrestrial organisms have evolved two solutions (Alpert and Oliver 2002). The first is to conserve water and avoid severe body water deprivation, such as the waxy coatings on shoots and the protective cocoon of the African lungfish *Protopterus annectens* (Procko and Shaham 2011). The second solution is to tolerate body water loss. A term, anhydrobiosis, is often used for the almost completely dehydrated but viable state of organisms facing extreme desiccation (Crowe, et al. 1992; Wharton 2015). Within either anhydrobiotic plants or animals, there is also a grade in desiccation tolerance. The most desiccation-tolerant organisms can enter the anhydrobiotic state and thus survive desiccation at any stage of their life cycle. The commonly studied organisms of this group are the bdelloid rotifers, the tardigrades, the resurrection plants (Farrant and Moore 2011; Costa, et al. 2017; Giarola, et al. 2017).

The DSBs induced by prolonged desiccation were suggested to open the “gateway to genetic exchange” and account for the elevated HGT frequency in bdelloid rotifers (Gladyshev, et al. 2008; Boschetti, et al. 2012; Flot, et al. 2013; Hespeels, et al. 2014; Eyres, et al. 2015; Debortoli, et al. 2016). However, the DSBs induced by prolonged desiccation in tardigrades (Neumann, et al. 2009) did not result in an elevated level of HGTs (Yoshida, et al. 2017). We first examine whether there are elevated levels of endosymbiotic DNA transfers (EDTs) in the desiccation-tolerant bdelloid rotifers and tardigrades. In the genomes of desiccation-tolerant bdelloid rotifers, *A. vaga* and *A. ricciae*, we detected 22 and 4 NUMTs, with total lengths of 32 kb and 14 kb, respectively. Similarly, in the genomes of desiccation-tolerant tardigrades, *H. dujardini* and *R. varieornatus*, we detected 14 and 53 NUMTs, with total lengths of 3.7 kb and 33 kb, respectively. By contrast, thousands of NUMTs were reported in the genome of honeybee *A. mellifera* (Pamilo, et al. 2007). We re-surveyed the honeybee genome with our parameters and detected 1791 NUMTs with a total length of 723.7 kb. Compared with a broad range of animals, the NUMT abundances of these four species are below the average value (Hazkani-Covo, et al. 2010; Hazkani-Covo and Martin 2017a). The astonishingly high level of HGTs in desiccation-tolerant bdelloid rotifers is not accompanied by an astonishing frequency of EDTs. On the other hand, in the honeybee that was claimed to have exceptionally high density of NUMTs (Pamilo, et al. 2007), so far as we know, there is no evidence for a high level of HGTs.

The resurrection plants can survive extreme drying and maintain a quiescent state for months to years (VanBuren 2017). To examine the relationship between prolonged desiccation and frequency of genetic exchange, we also surveyed the abundance of NUPTs in the resurrection plants. In the flowering plant *D. hygrometricum* and the spike moss *S. tamariscina*, we detected 1610 NUPTs with a total length 467 kb and 656 NUPTs with a total length of 521 kb, respectively. The abundance of NUPTs in these resurrection plants is much higher than that of NUMTs in desiccation-tolerant bdelloid rotifers and tardigrades. However, previous studies have shown that the plant genomes generally have much higher NUMT/NUPT contents than animals (Hazkani-Covo, et al. 2010; Hazkani-Covo and Martin 2017a). The NUPT contents we observed in resurrection plants are not excessive in plants.

### Comparison of NUMT/NUPT Contents between Organisms Differing in Desiccation-Tolerance

In the second rank of desiccation-tolerance, anhydrobiosis is restricted to particular developmental stages, like the dormant eggs water flea *Daphnia*, the cysts of brine shrimp, *Artemia*, and the orthodox seeds of most angiosperms (Farnsworth 2000; Gusev and Okuda 2012). In the last rank, all the stages of their life cycle are desiccation sensitive, including the eggs of animals and the seeds of viviparous plants. From this aspect, a viviparous plant, like *F. esculentum*, is more sensitive to desiccation than a plant with orthodox seeds, like *C. quinoa* (Carjuzaa, et al. 2008; Pirzadah, et al. 2016).

To approach a generally conclusion on the relationship between desiccation and frequency of genetic exchange, we compared nine pairs of organisms within each pair the organisms are different in their desiccation-tolerance (Table 1). The most closed relatives used as controls of the desiccation-tolerant organisms were selected referring widely used phylogenetic databases including Timetree (http://timetree.org), NCBI taxonomy (https://www.ncbi.nlm.nih.gov/taxonomy), and Angiosperm Phylogeny Website (http://www.mobot.org/MOBOT/research/APweb/welcome.html) (Gabriel, et al. 2007; Federhen 2012; Kumar, et al. 2017). In cases where one phylogenetic branch contains two desiccation-tolerant species or two desiccation-sensitive species, we used the average value of the two species. For example, the average value of desiccation-tolerant bdelloid rotifers, *A. vaga* and *A. ricciae*, was used to compare the average value of desiccation-sensitive bdelloid rotifers, *R. magnacaicarata* and *R. macrura*. Nonparametric pairwise comparison did not detect a significant difference between desiccation-tolerant species and their desiccation-sensitive relatives in either the number of NUMTs/NUPTs or the total length of NUMTs/NUPTs (Wilcoxon Signed ranks test, *P* > 0.05, fig. 1). A large nuclear genome is expected to have more sites for NUMT/NUPT integration and thus likely to have more NUMTs/NUPTs. To control the influence of nuclear genome size, we compared the density of NUMTs/NUPTs in nuclear genomes. Still, no significant differences were detected between desiccation-tolerant species and their desiccation-sensitive relatives (Wilcoxon Signed ranks test, *P* > 0.10, fig. 2).

**Table 1.**
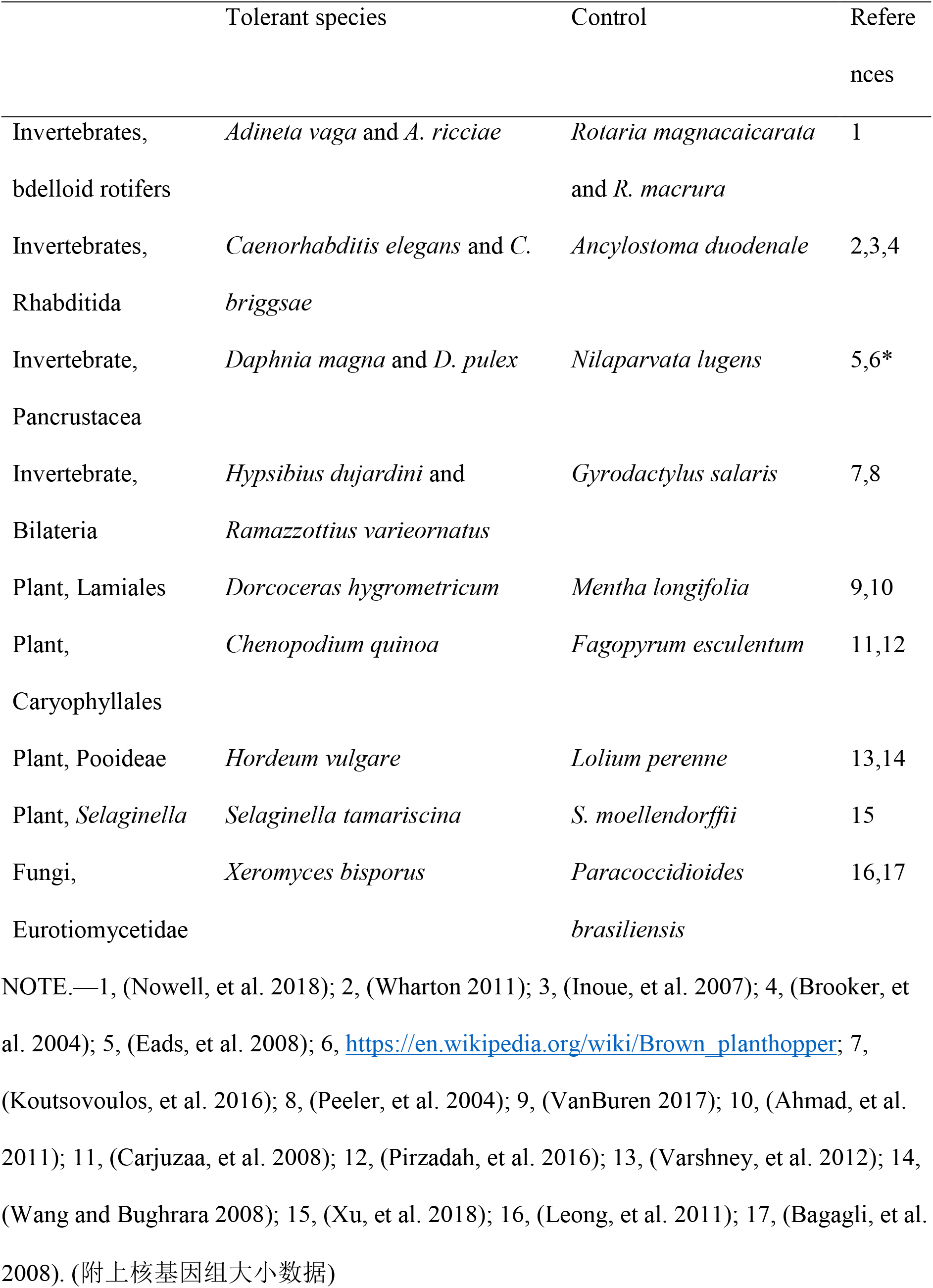
Desiccation-Tolerant Organisms and Their Desiccation-Sensitive Relatives Used in This Study.

**Fig. 1.**
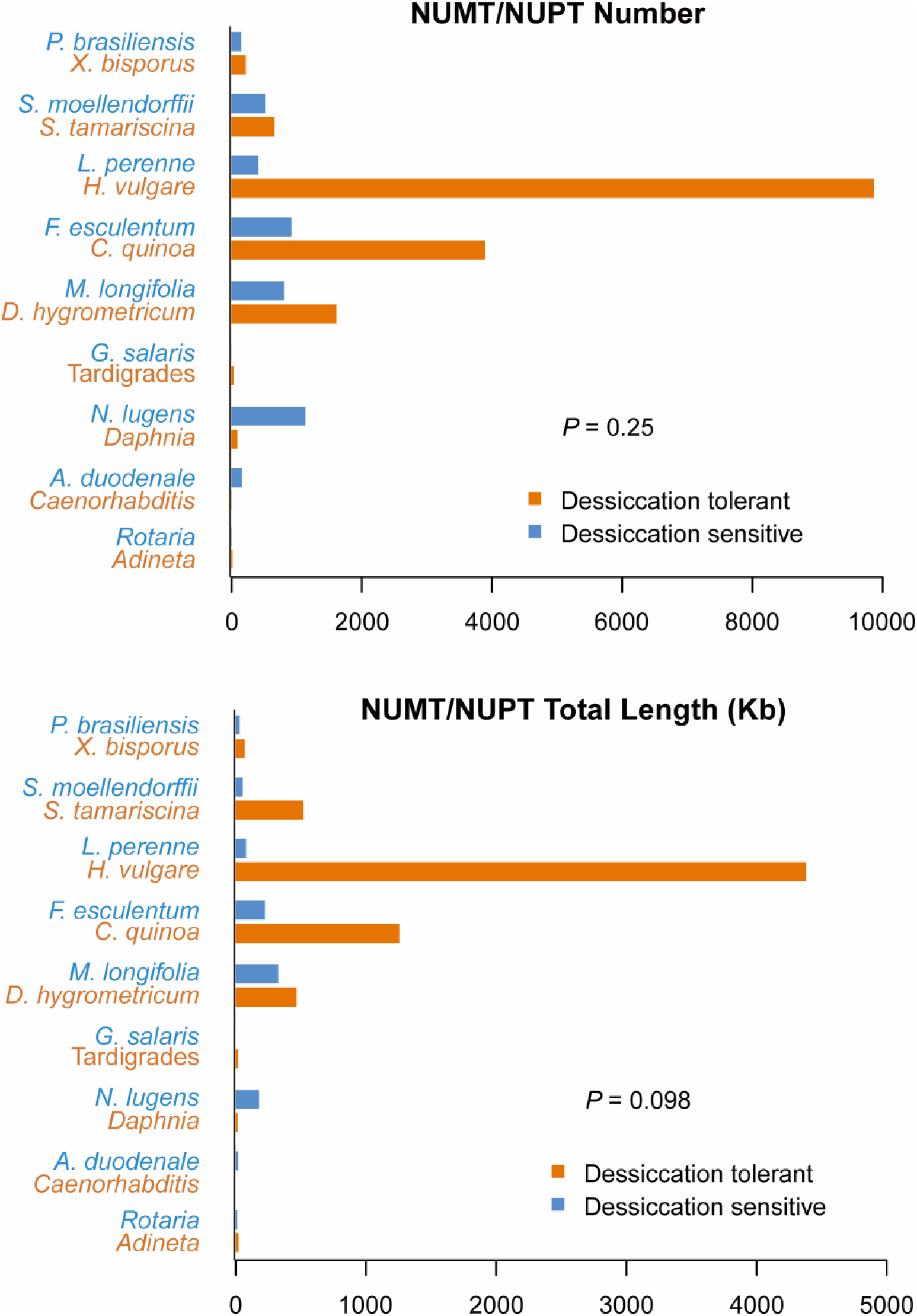
Comparison of the NUMT/NUPT contents between desiccation-tolerant organisms and their desiccation-sensitive relatives. The average value of *Adineta vaga* and *A. ricciae* was presented in this figure and marked by their genus name, *Adineta*. It is the same for *Rotaria magnacaicarata* and *R. macrura*, *Daphnia magna* and *D. pulex*, and *Caenorhabditis elegans* and *C. briggsae*. The average value of *Hypsibius dujardini* and *Ramazzottius varieornatus* was presented in this figure and marked by their common name, tardigrades. Wilcoxon Signed Ranks tests (two tailed) were used to calculate the *P* value.

**Fig. 2.**
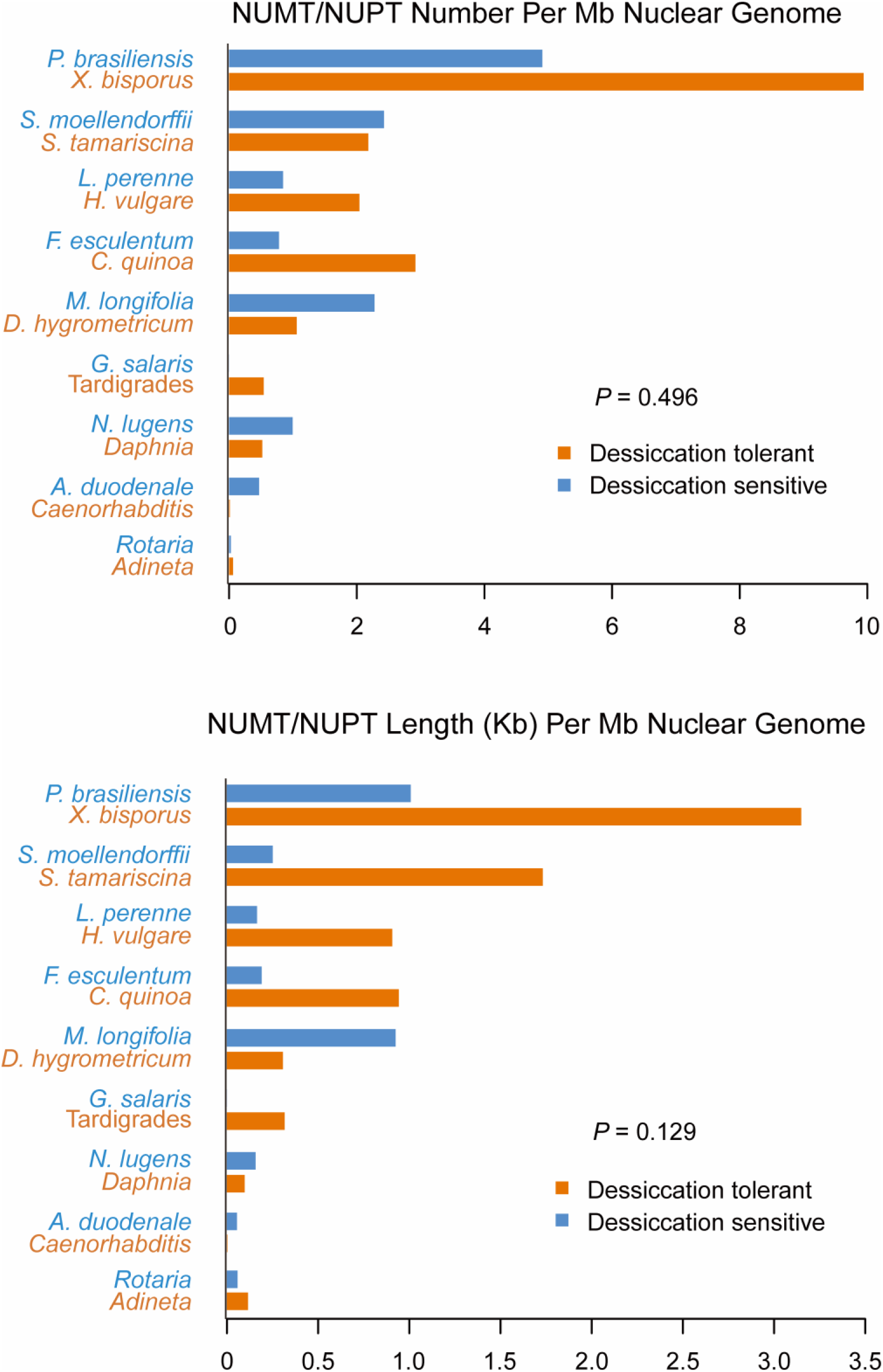
Comparison of the NUMT/NUPT densities between desiccation-tolerant organisms and their desiccation-sensitive relatives. The average value of *Adineta vaga* and *A. ricciae* was presented in this figure and marked by their genus name, Adineta. It is the same for *Rotaria magnacaicarata* and *R. macrura*, *Daphnia magna* and *D. pulex*, and *Caenorhabditis elegans* and *C. briggsae*. The average value of *Hypsibius dujardini* and *Ramazzottius varieornatus* was presented in this figure and marked by their common name, tardigrades. Wilcoxon Signed Ranks tests (two tailed) were used to calculate the *P* value.

Although most of the EDTs are by direct transfer of organelle DNA into the nucleus, there is also evidence supporting EDTs mediated by RNA molecules (Timmis 2012). Therefore, we also compared the NUMTs/NUPTs contributed by the transcribed regions of organelle genomes, i.e., protein-coding genes, tRNA, and rRNA. Similar to the above results, no significant difference was detected between desiccation-tolerant species and their desiccation-sensitive relatives (Wilcoxon Signed ranks test, *P* > 0.10). Although both nuclear genomes and mitochondrial genomes of *H. vulgare* and *L. perenne* are available for analysis, we just includes their NUMTs contributed by transcribed sequences in our comparison, but not the NUMTs contributed by the full mitochondrial genome sequences. That is because nuclear sequences are occasionally transferred to mitochondrial genomes in plants (Goremykin, et al. 2012; Mower, et al. 2012). It is very difficult to determine the direction of transfer when an uncharacterized sequence is present in both the mitochondrial genome and the nuclear genome.

## Discussion

Desiccation is a widely observed cause of DSB formation from bacteria, plants to animals such as tardigrades and bdelloid rotifers (Mattimore and Battista 1996; Faria, et al. 2005; Pitcher, et al. 2007; Neumann, et al. 2009; Gusev, et al. 2010; Waterworth, et al. 2011; Gusev and Okuda 2012; Hespeels, et al. 2014; Ricci 2017). Organisms that frequently encounter desiccation stress have evolved efficient mechanisms to repair the DSBs (de Groot, et al. 2009). Accordingly, the radioresistance of *Deinococcus radiodurans* and bdelloid rotifers has been attributed to a consequence of their adaptation to desiccation stress (Mattimore and Battista 1996; Gladyshev and Meselson 2008). The DSBs in nuclear genomes are opportunities for foreign DNA integration and so was postulated as the gateway for HGTs (Hespeels, et al. 2014). If the DSBs of the desiccation-tolerant organisms are efficient gateway for foreign DNA integration, the frequencies of EDTs should also be dramatically elevated. However, we did not observed excessive NUMTs/ NUPTs in the genomes of extreme desiccation-tolerant organisms, bdelloid rotifers *A. vaga* and *A. ricciae*, tardigrades, *H. dujardini* and *R. varieornatus*, or the resurrection plants *D. hygrometricum* and *S. tamariscina*. Further statistically comparison of nine pairs of desiccation-tolerant organisms with their desiccation-sensitive relatives did not find evidence for desiccation-elevated EDT frequency, either.

The induction of DSBs by desiccation is a consensus. But frequent DSBs do not necessarily increase the frequency of foreign DNA integration into the nuclear genomes. The integrations of foreign DNA, i.e., HGTs and EDTs, depend on the specific mechanisms of DSB repair, which fall into two main categories: homologous recombination (HR) and non-homologous end joining’ (NHEJ). Capture of foreign DNA sequences is possible only when DSBs are repaired by a subtype of NHEJ, named the alternative end joining (alt-EJ), which is much less frequently used than other mechanisms (Ono, et al. 2015; Ceccaldi, et al. 2016). The choice of repair pathway between HR and NHEJ and among different subtypes of NHEJ depends on phylogenetic positions, cell cycle phases and chromosomal repetitive contents (Lieber and Karanjawala 2004; Mao, et al. 2008; Her and Bunting 2018). If an organism predominantly use HR in its DSB repair, desiccation-induced DSBs do not open the gateway for HGTs and EDTs. In general, organism with compact genomes like yeasts preferentially use HR while in mammals and plants NHEJ is the predominant DSB repair pathway. This might explain the generally higher NUMT/NUPT contents in flowering plants previously reported as well as we observed (fig. 1).

If the desiccation-induced DSBs open the gateway for foreign DNA, but the donor is not efficient, the frequency of NUMT/NUPT could not be elevated by desiccation. Evidence for the effects of the availability of foreign DNA on the frequency of transfer came from analysis of the organisms that have single plastid or mitochondrion in each cell. Lysis of these organelles would almost certainly result in death of the cells. These organisms, with the limited availability of donor organelle DNA, have been shown to have much lower amounts of NUMTs or NUPTs than organisms with multiple organelles per cell (Smith, et al. 2011). All the organisms we studied have multiple organelles per cell. Instead, the organelles of desiccation-tolerant organisms seems to be more efficient DNA donors because of organelle membrane disruption during desiccation.

As insertion mutations, the abundance of NUMTs/NUPTs as well as the frequency of HGT depend not only on its insertion rate but also on the fixation possibility in evolution (Schonknecht, et al. 2014). For a compact nuclear genome with a limited amount of junk DNA sequences, most insertions of NUMTs/NUPTs would disrupt the nuclear genetic information and so be selected against. Similar to (Hazkani-Covo, et al. 2010; Smith, et al. 2011; Yoshida, et al. 2014), we observed a significant positive correlation between the abundance of NUMTs/NUPTs and nuclear genome size in our dataset (Spearman’s rho = 0.63 and 0.60 for NUMTs/NUPTs number and total length, respectively, and *P* < 0.05 for both cases).

Finally, it should be noted that the low level of EDTs in bdelloid rotifers *A. vaga* and *A. ricciae* should not be regarded as evidence against the reliability of previous reports on the elevated levels of HGTs in bdelloid rotifers. Our results are just inconsistent with the hypothesis that desiccation could facilitate gene transfer by inducing DSBs in nuclear genomes. The bdelloid rotifers might have elevated levels of HGTs unrelated with desiccation. Actually, Nowell et al. 2018 recently found high levels of HGTs in both desiccation-tolerant and desiccation-sensitive bdelloid species (Nowell, et al. 2018). In addition, our statistically analysis was based on a very small sample. With the accumulation of sequenced nuclear genomes and organelle genomes, it is necessary to compare the desiccation-tolerant and desiccation-sensitive species in a large scale, especially in lineages that have large nuclear genomes whose DSBs are repaired predominantly by NHEJ.

## Acknowledgments

This work was supported by the National Natural Science Foundation of China (grant numbers 31671321).

